# Distinct neural states encode task identity in frontal eye field and interact with its core spatial properties

**DOI:** 10.1101/2023.09.05.556340

**Authors:** A. Mouille, C. Gaillard, E. Astrand, C. Wardak, S. Ben Hamed, J.L. Amengual

## Abstract

The prefrontal cortex (PFC) plays a key role in selecting, maintaining and representing sensory information, and in its integration with our current internal goals and expectations, implementing such cognitive processes as executive functions, attention, decision making and working memory. Performing computation over all of these functional cognitive processes and dynamically shifting from one to the other based on task demands requires a complex functional organization and a high degree of coding flexibility. The PFC cells show a non-linear mixed selectivity characterized by specific tuning for multiple task- and behaviour related parameters. This non-linear mixed selectivity thought to allow for a high-dimensional representation of information. Here, we asked if the PFC is mainly involved in specific task-parameters representation, or if it additionally holds a higher order representation for task-identity. We thus trained two macaques to perform three different tasks: a memory guided saccade task and two detection tasks involving different attention mechanisms. Multi-unit activity was recorded in the frontal eye field, bilaterally, while monkeys performed these three tasks in a same session. Using demixed Principal Component Analysis, we found a two-dimensional neural state that characterized each of these tasks. The lower dimensional representation of the activity recorded during the performance of the two attentional tasks were more similar to each other than to the memory-guided saccade task. Furthermore, we report that task and spatial information are non-linearly mixed, a signature of a high-dimensional neural representation. Overall, this indicates that PFC encodes task identity information and flexibly adjusts its sensory processes as a function of the specific ongoing task.

**Significance Statement:** One long lasting question in cognitive neuroscience is whether PFC mainly represents the different parameters that are needed to perform any given task, or if it additionally holds a higher order representation for task-identity. We recorded from the macaque frontal eye fields while monkeys performed a memory guided saccade task and two detection tasks involving different attention mechanisms. We found a task-identity neural state in which the two attentional tasks were represented similarly to each other but differently from the memory-guided saccade task. We also report that task and spatial information are non-linearly mixed, a signature of a high-dimensional neuronal representation. Overall, this indicates that PFC encodes task identity information and flexibly adjusts its sensory processes as a function of the specific ongoing task.

## Introduction

The prefrontal cortex (PFC) produces internal representations of our actions and of our environment. To do so, PFC neural computations contribute to the selection, maintenance and filtering of sensory information, in order to integrate the representation of the current task parameters with overarching internal goals and expectations. In doing so, PFC functionally contributes to several cognitive processes that include executive functions (Friedman and Robbins, 2022; Goldman-Rakic, 1996; Robbins, 1996), attention (Amengual and Ben Hamed, 2021; Ibos et al., 2013; Paneri and Gregoriou, 2017; Rossi et al., 2009), decision making (Domenech and Koechlin, 2015; Funahashi, 2017) and working memory (Funahashi and Kubota, 1994; Lara and Wallis, 2015). Such rich functional architecture is essential to guide the cognitive control of our actions and to shape our behaviour. This vast mosaic-like map of functions demands a very fine functional organization that allows performing multiple computations minimizing interference. Electrophysiological studies have shown that PFC displays cells tuned to multiple task- and behaviour-related parameters, a property called non-linear mixed selectivity, which provides computational advantages for information transmission to downstream neurons (Fusi et al., 2016; Rigotti et al., 2013). In the present study, we aim to describe how this mixed-selectivity allows the PFC to hold a representation of the ongoing task in addition to a representation of sensory information that dynamically changes as a function of specific task demands.

Non-linear mixed-selectivity has been proposed to result in a high-dimensional representation of multiple sensory information patterns of stimuli (such as position, colour or shape) (Rigotti et al., 2013). Such encoding strategy allows simple readout of complex patterns of behaviour, using for example linear classifiers. One important question is whether non-linear mixed selectivity cells in the PFC are merely involved in the representation of specific task-parameters such as position, color or attention irrespective of the ongoing task, or whether these cells represent higher cognitive levels of encoding by additionally representing task-identity, as a proxy for the implementation of a specific cognitive process (e.g. attention versus spatial working memory). If task-identity is represented, is this information represented at the same hierarchical level as specific task parameters (i.e. independently), or is task identity represented at a higher hierarchical level (i.e. task information changing for example how the spatial location of stimuli is represented)?

To address this question, we trained two macaque monkeys to perform three different tasks in the same recording sessions: 1) a centrally cued manual target detection task, the target being presented in the periphery of the visual field; 2) a cued manual target detection task, in which the cue is presented at the expected location of the target in the periphery of the visual field; 3) a memory guided saccade task in which an eye movement has to be performed, upon instruction, towards a previously cued location. Each of these tasks recruited specific cognitive functions: endogenous attention, exogenous attention and spatial working memory. While monkeys were performing these tasks in independent blocks or trials, dense recording probes were used to record multi-unit activity (MUA) from both frontal eye fields (FEF), a region in the prefrontal cortex linked with top-down attentional processes (Astrand et al., 2016; Bruce et al., 1985; Goldberg and Segraves, 1987; Ibos et al., 2013) and working memory (Noudoost et al., 2021; Offen et al., 2010). Using a demixed principal component analysis (dPCA (Kobak et al., 2016; Machens, 2010)), we extracted latent components associated specifically with either sensory information or task identity. We found that the neuronal processes underlying each task were encoded in two different orthogonal components, one component reflecting the encoding of either attention versus working memory, and the other component reflecting the encoding of endogenous versus exogenous attentional enhancement. In addition, we identified a component that encoded the interaction between task-identity and sensory processing, indicating that these two sources of information (task and position of the sensory information) could be decoded simultaneously by a single linear readout, though the readout of sensory information varied across tasks.

All in all, these results suggest that prefrontal cortex encodes sensory information and task-identity in a high-dimensional neural representation. In addition, spatial information is represented independently from task-related information, indicating that task-identity is represented at the same hierarchical level as specific task parameters such as position. At the fundamental level, it will be important to pursue on this work in order to better understand the specific dimensions that the prefrontal cortex uses to represent task identity. At the translational level, this work opens new venues for the implementation of brain computer interface technologies based on machine learning algorithms able to extract complex cognitive information from real-time PFC recordings.

## Methods

### Surgical procedure and FEF mapping

Two male rhesus monkeys (Macaca mulatta) weighing between 6-8 kg underwent a unique surgery during which they were implanted with two MRI compatible PEEK recording chambers placed over the left and the right FEF hemispheres respectively, as well as a head fixation post. Gas anaesthesia was carried out using Vet-Flurane, 0.5 – 2% (Isofluranum 100% at 1000 mg/g) following an induction with Zolétil 100 (Tiletamine at 50mg/ml, 15mg/kg and Zolazepam, at 50 mg/ml, 15 mg/kg). Post-surgery pain was controlled with a morphine painkiller (Buprecare, buprenorphine at 0.3mg/ml, 0.01mg/kg), 3 injections at 6 hours’ interval (first injection at the beginning of the surgery) and a full antibiotic coverage was provided with Baytril 5% (a long action large spectrum antibiotic, Enrofloxacin 0.5mg/ml) at 2.5 mg/kg, one injection during the surgery and thereafter one each day during 10 days. A 0.6mm isomorphic anatomical MRI scan was acquired post surgically on a 1.5T Siemens Sonata MRI scanner, while a high-contrast oil-filled 1mmx1mm grid was placed in each recording chamber, in the same orientation as the final recording grid. This allowed a precise localization of the arcuate sulcus and surrounding grey matter underneath each of the recording chambers. The FEF was defined as the anterior bank of the arcuate sulcus and we specifically targeted those sites in which a significant visual, premotor and/or oculomotor activity was observed during a memory guided saccade task at 10 to 15° of eccentricity from the fixation point. In order to maximize task-related neuronal information at each of the 24-contacts of the recording probe, we only recorded from sites with task-related activity observed continuously over at least 3 mm of depth. All surgical and experimental procedures were approved by the local animal care committee (C2EA42-13-02-0401-01) approved by the French Ministry of Research and in compliance with the European Community Council, Directive 2010/63/UE on Animal Care.

### Behavioural tasks and experimental setup

In all behavioural tasks, monkeys were seated in front of a computer screen (1920x1200 pixels and a refresh rate of 60 Hz) at a distance of 45 cm with their head fixed. To initiate a trial, the monkeys had to hold a bar in front of the animal chair, thus interrupting an infrared beam. They were required to hold the bar throughout the trial until a target stimulus was presented, otherwise the trial was aborted. The eccentricity of the visual stimuli was adjusted from day to day between 10 to 15°, to match the preferred spatial location of the multi-unit activity recorded on the 48 contacts of both recording probes.

### Exogenous 100% validity cued luminance change detection task (*Exo*)

The task is a 100% validity cued luminance change detection task with temporal distractors (Figure 1A). To initiate a trial, the monkeys had to hold a bar in front of the animal chair, thus interrupting an infra-red beam. A blue fixation cross (size: 0.7×0.7°) appeared in the centre of the screen. Monkeys were required to hold fixation throughout the entire trial, within a fixation window of size 4°x4°. Failing to do so aborted the trial and another trial started again. Four grey landmarks (0.5×0.5° for monkey M1, 0.68×0.68° for monkey M2) were presented simultaneously and equidistantly with the fixation cross. These landmarks were located, in the upper right, upper left, lower left and lower right quadrants of the screen, thus defining the corners of an illusionary square. Their specific eccentricity was adjusted from day to day between 10 to 15°, to match the preferred spatial location of the multi-unit activity recorded on the 48 contacts of both recording probes. After a variable delay from fixation onset, ranging between 700 and 1900 ms, a green square was presented for 350 ms, indicating to the monkey in which of the four landmarks the rewarding target change in luminosity would appear. This green square (from now on, the cue) was presented on top of the peripheral landmark to be attended to. After the cue presentation, the monkeys needed to orient their attention to the target landmark in order to monitor it for a change in luminosity while maintaining eye fixation onto the central blue cross. This change of luminosity (from now on, the target) could occur anywhere between 500 to 2800 ms from cue onset. In order to receive their water or juice reward, the monkeys were required to release the bar (thus restoring the infra-red beam) in a time window of 200 to 700 ms following the target onset. This event accounted as a hit trial, while failing to respond to the target resulted in miss trials. In order to assure that the monkeys were correctly orienting their attention towards the cued landmark, unpredictable changes in the luminosity identical to the awaited target luminosity change could take place at the uncued landmarks (from now on, distractors). On each trial, from none to three such unpredictable distractors could take place, no more than one per uncued landmark position. Monkeys were trained to avoid responses to these distractors as its response interrupted the trial and was counted as a false alarm trial.

**Figure 1.**
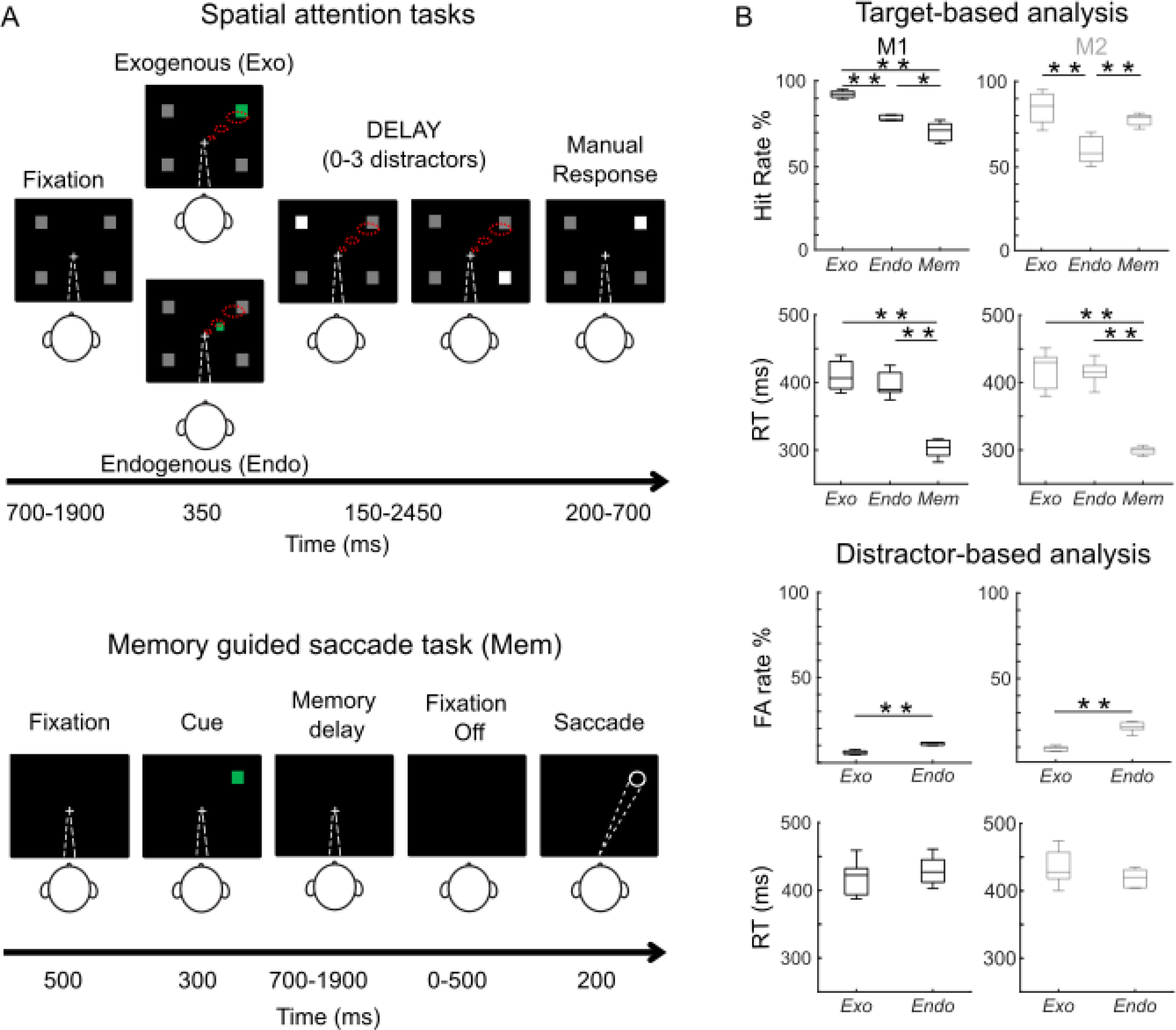
Description of the tasks and main behavioural results. (A) Monkeys performed three behavioural tasks: The endogenous (*Endo*) and exogenous (*Exo*) attention task consisted of 100 % validity cued target-detection task with distractors. To initiate the trial, monkeys had to hold a bar with the hand and fixate their gaze (white dashed line, first screen) on a central cross on the screen during the whole trial. A cue (green square, second screen) was presented indicating to the monkey in which of the four landmarks the target change in luminosity would appear. Monkeys received a liquid reward for releasing the bar 200-700 ms after target presentation onset. Monkeys had to ignore any un-cued event (distractors). Importantly, they had to detect the stimuli using selective spatial attention (red dashed lines). In the memory-guided saccade task (Mem), a peripheral stimulus appeared 300ms following fixation (500 ms). The monkeys had to hold the position in memory, throughout a variable delay of 700 to 1900ms, until the fixation cross disappeared. At the extinction of the fixation cross, the monkeys had 500 ms to execute a saccade to the previously presented stimulus location. At the end of the saccade, the peripheral stimulus reappeared and the monkeys had to wait and respond when a target stimulus was presented. (B) Behavioural performance. On the top part, boxplots represent the distributions of hit rates and reaction times (RT) over all sessions (n=12). On the bottom part, boxplots show distributions of the false alarm rate (FA rate) and reaction time (RT) of responses to distractors over all sessions (n=12, non-parametric Wilcoxon signed rank test *p<0.05, **p < 0.01).

### Endogenous 100% validity cued luminance change detection task (*Endo*)

The *endogenous* task was identical to the exogenous task in all aspects except for the location and size of the cue (Fig. 1B, panel 1). Cue stimulus consisted in a green square (0.2×0.2° for monkey M1 and 0.3×0.3° for monkey M2) and it was presented close to the fixation cross in the same direction as the landmark to be attended (at 0.3° for monkey M1 and at 1.1° for monkey M2, from the fixation point).

### Memory-guided saccade task (*Mem Sacc*)

A standard oculomotor memory task was used (see fig. 1B, panel 3) in which the monkeys needed to fixate a blue cross in the centre of the screen (0.3×0.3°) in order for the trial to be initiated. Fixation within a window of 4×4° around the fixation cross was required or else the trial was aborted. 500 ms following fixation, a peripheral stimulus (size: 0.3×0.3°) appeared for 300 ms. The monkeys had to hold the position in memory, throughout a variable delay of 700 to 1900 ms, until the fixation cross disappeared. At the extinction of the fixation cross, the monkeys were required to execute a saccade to the previously presented peripheral stimulus, within a window of 8×8° around it. The monkeys had 500 ms to execute the saccade to the correct position or the trial was aborted. At the end of a correct saccade, fixation on the empty screen had to be maintained 200 ms before the peripheral stimulus reappeared. The monkeys then had to fixate this peripheral stimulus within a window of 4×4° and wait until the red colour changed to grey (variable delay of 500 to 800 ms). At colour change, the monkeys had to release the bar (thus restoring the infrared beam) within a time window of 150 to 800 ms for a liquid reward.

Within any recording session, the monkeys performed the three tasks consecutively. The order of the attentional tasks was changed randomly between sessions, while the memory-guided saccade task was always presented in the third order, as monkeys displayed a strong behavioural preference for this task.

### Neural recordings

Simultaneous recordings in the two FEF/hemispheres were carried out using two 24-contact Plexon U-probes. The contacts had an interspacing distance of 250 µm. Neural data was acquired with PlexonOmniplex® neuronal data acquisition system (Omniplex). The data was amplified 400 times and digitized at 40,000 Hz. The neuronal data was low-cut filtered at 300 Hz. All analyses were performed on the MUA recorded from each of the 48 recording contacts. A threshold defining the MUA was applied independently for each recording contact and before the actual task-related recordings started. All further analyses of the data were performed in Matlab (The Mathworks INC., Natick, Massachussetts).

### Data pre-processing

In each task, MUA activity was epoched per trial. Spike trains were filtered with a Gaussian kernel (δ = 60 ms). First, MUA activity previous to the cue onset was epoched from -500 ms to 0 with respect to the onset of the Cue for the *Pre-Cue analysis*. Then, MUA activity was epoched between 500 ms to 0 ms prior to Target onset for the *Delay analysis*. To extract the averaged firing rates, these epoched MUA data were baselinecorrected (independently for each channel and each trial) relative to the time period from -500 ms to -300 ms before the cue.

### MUA selectivity

Spatially selective channels were identified using a non-parametric Wilcoxon test of the activity between the baseline and a post cue period (300 to 600 ms following cue onset). For each spatially selective channel, the most and least preferred positions were further identified and respectively corresponded to the channels with the higher and lower firing rates among the four possible cued locations measured in the time interval 300 to 600 ms following cue onset.

### Demixed PCA analysis

Recent studies have shown that neurons in the prefrontal cortex show simultaneous significant tuning to multiple task related parameters, a property called mixed selectivity(Rigotti et al., 2013). This feature is a hallmark of high dimensionality of the neuronal population. Due to this property, the activity structure of the population can be estimated by applying a dimensionality reduction method to the recorded activity such as Principal Component Analysis (PCA). Using this method, it is possible to extract a number of latent variables (principal components) that capture independent sources of data variance providing a description of the statistical features of interest (Cunningham and Yu, 2014). However, PCA method is blind to the sources of variability of the data and does not take into account the task related parameters, mixing these sources of information within each of the extracted latent variables (Kobak et al., 2016). Here, we wanted to study and compare the population activity structure in the three different tasks and describe how much the variance in the neuronal population can be explained by task-related aspects such as task, spatial target location, and their interaction. To address this, we applied dPCA (demixed Principal Component Analysis (Kobak et al., 2016; Machens, 2010)) on the data which captures a maximum amount of variance explained by predefined sources of variability in each extracted component and reconstruct the time course of the category-specific response modulation.

Demixed PCA was applied in the two different time intervals (*Pre-Cue* and *Delay*). The purpose of the first dPCA analysis was to identify specific task-related components during the Pre-Cue period. Then, another dPCA was performed during the delay period, with the aim to extract task and spatial-related components, as well as components associated with their interaction (non-linear mixed selectivity). Both analyses were performed using appropriate Matlab based toolbox developed by Kobak and colleagues(Kobak et al., 2016). We considered the 20 first demixed components that accounted for > 90% of the total variance.

We used the decoding axis of each dPC (assigned to task in the *Pre-Cue* analysis, or task, position and interaction in the *Delay* analysis) as a linear classifier to decode the different types of trials (*Endo, Exo* and *Mem Sacc* for task, and *Up Left, Up Right, Down Left* and *Down Right* for position), following the procedure described by Kobak and colleagues (Kobak et al., 2016). This method allows to quantify the capacity of each demixed component to classify a trial between the classes of a given category. To extract the statistical significance of this accuracy, we shuffled 100 times all available trials between classes and we thereby computed the distribution of classification accuracies expected by chance. For each session, the chance-level was considered as the maximal accuracy value obtained across all randomizations.

## Results

### Monkey performance is task-dependent

The present study aims to describe and compare, in the PFC, the electrophysiological underpinnings of spatial information encoding during the performance of different tasks. To this end, we recorded the MUA from bilateral monkey frontal eye field (FEF), an area involved in the control of covert spatial attention (Armstrong et al., 2009; Gregoriou et al., 2009; Ibos et al., 2013; Paneri and Gregoriou, 2017) and saccades (Hanes et al., 1995; Schall, 2004, 1991; Vernet et al., 2014), of two different macaque rhesus monkeys while they performed three different tasks: an endogenous attention task (*Endo*), an exogenous attention task (*Exo*) and a memory-guided saccade task (*Mem*). These three tasks were presented consecutively in the same session in a block design. The schema of these tasks was very similar (see Figure 1A and Methods section for an extensive description of these tasks): at the beginning of each trial, a visual cue was presented in one of the four possible screen quadrants, and after a variable and randomized time interval monkeys had to produce a response. However, the actual task differed and the monkeys needed to adapt their cognitive processes to perform the tasks. The exogenous and endogenous attentional tasks consisted in a 100% validity cued luminance change detection task in which monkeys had to respond manually to a target located peripherally in the cued quadrant. In the exogenous version, the cue was shown in the peripheral landmark to be attended to (thus, in the same position as the target). In contrast, in the endogenous attention task the cue was presented close to the fixation cross pointing towards the landmark to be attended. In both endogenous and exogenous attention tasks, distractor stimuli were presented during the cue to target interval (CTOA), and monkeys needed to avoid them (otherwise, the trial was cancelled). These two tasks were used to recruit attentional processes (Astrand et al., 2016). In the memory-guided saccade task, monkeys were required to hold the position of a spatial cue in memory during a variable amount of time and, then, to perform a saccade towards that memorized spatial location on the presentation of a go signal. Importantly, and in contrast with the attentional tasks, monkeys were required to perform a spatially oriented oculomotor response rather than a manual non-oriented response. This task thus recruited spatial working memory processes. Although these three tasks showed some similarities in their structure, they recruited different cognitive processes. While monkeys needed to enhance visual attention in the cued quadrant to respond to the target onset in the exogenous and endogenous attentional tasks (Amengual et al., 2022; Astrand et al., 2020, 2016), they needed to memorize the location of the cue to produce an oriented eye movement towards that location.

Both monkeys showed different behavioural patterns as a function of the type of task they performed. Non-parametric Friedman test revealed that hit rates (M1: χ^2^(5) = 12.29, p = 0.002; M2: χ^2^(5) = 7.6, p = 0.0224) and reaction times (RT) (M1: χ^2^(5) = 12.29, p = 0.0021; M2: χ^2^(5) = 7.6, p = 0.0224) were different across tasks (Figure 1B). This reflects different task demands.

Both monkeys were able to discriminate the change in luminosity of the cued landmark while ignoring distractors in the spatial attention tasks, though their performance differed depending on the type of task. Indeed, both monkeys showed a higher hit rate when performing *Exo* than *Endo*, (fig.1.B M1: 92% vs. 78%, Wilcoxon ranksum test, Z =3.07, p=5.8e-04; M2: 87% vs. 60%, Wilcoxon ranksum test, Z=2.09, p=0.0079). M1 showed a lower hit rate in memory-guided saccade task than in *Exo* and *Endo*, while M2 showed significantly lower hit rate in *Endo* than in *Mem* (Figure 1B M1: 72% vs. 92%, Wilcoxon ranksum test, Z =3.07, p=5.8e-04, 72% vs.78%, Z =2.17, p=0.0233; M2: 60% vs. 78%, Z=2.51 p=0.0079). Regarding the RT, and expectedly, both monkeys had slower manual response times in attentional tasks than saccadic response times in the *Mem* task (M1, Friedman test, χ^2^(5) =12.29, p = 0.0021; M2, Friedman test, χ^2^(5) = 7.6, p = 0.022. M1: *Exo* vs *Mem*, Wilcoxon ranksum test, Z=3.07, p=5.8e-4, *Endo vs Mem*, Wilcoxon ranksum test, Z=3.07, p=5.8e-4; M2, *Exo vs Mem*, Wilcoxon ranksum test, Z=2.51, p=0.0079; *Endo vs Mem*, Z=2.51, p=0.0079). Regarding the attentional tasks, in despite of no significant difference, RTs tended to be slower for the Endo than for the *Exo task* as reported in other studies (Astrand et al., 2016).

False alarm rates (measured as the ratio between the number of responses to distractor and the total number of distractors shown) and reaction time in false alarms were analysed only for the attentional tasks, since no distractors were used in *Mem*. Both monkeys showed lower false alarm rates in *Endo* than in *Exo* (fig.1.B M1: 7% vs. 13%, Wilcoxon ranksum test, M1: Z=2.56 p=0.007; M2: 10% vs. 21%, Wilcoxon ranksum test, Z=0.63, p=0.0079). Monkeys did not show differences in reaction times in the false alarms between the two attentional tasks. To note, responses were slower in false alarms than in hits for M1 in *Endo* (Wilcoxon ranksum test, Z=1,79, p=0.0175).

### Spiking activity in the FEF encodes task- and sensory-related information simultaneously

We examined the FEF neural responses during the performance of these three tasks. To this end, we averaged the smoothed firing rates in the time interval -500 ms to 1000 ms locked to the cue for each of the three tasks. As reported in previous studies (Amengual et al., 2022; Di Bello et al., 2021; Ibos et al., 2013; Moore and Armstrong, 2003), these recorded neurons show enhanced responses when the cue was presented in or was cueing towards their preferred spatial position, relative to when it was presented in the least preferred spatial location, and this for all three tasks (Figure 2A&B). Similar results were found when data was locked to the target onset (Supplementary Figure 1). Spatial selectivity however varied between the three tasks. Indeed, we found a significant task-effect on the response to the preferred position (non-parametric Friedman test χ^2^(5) = 39.3, p = 2.89e-09).

**Figure 2.**
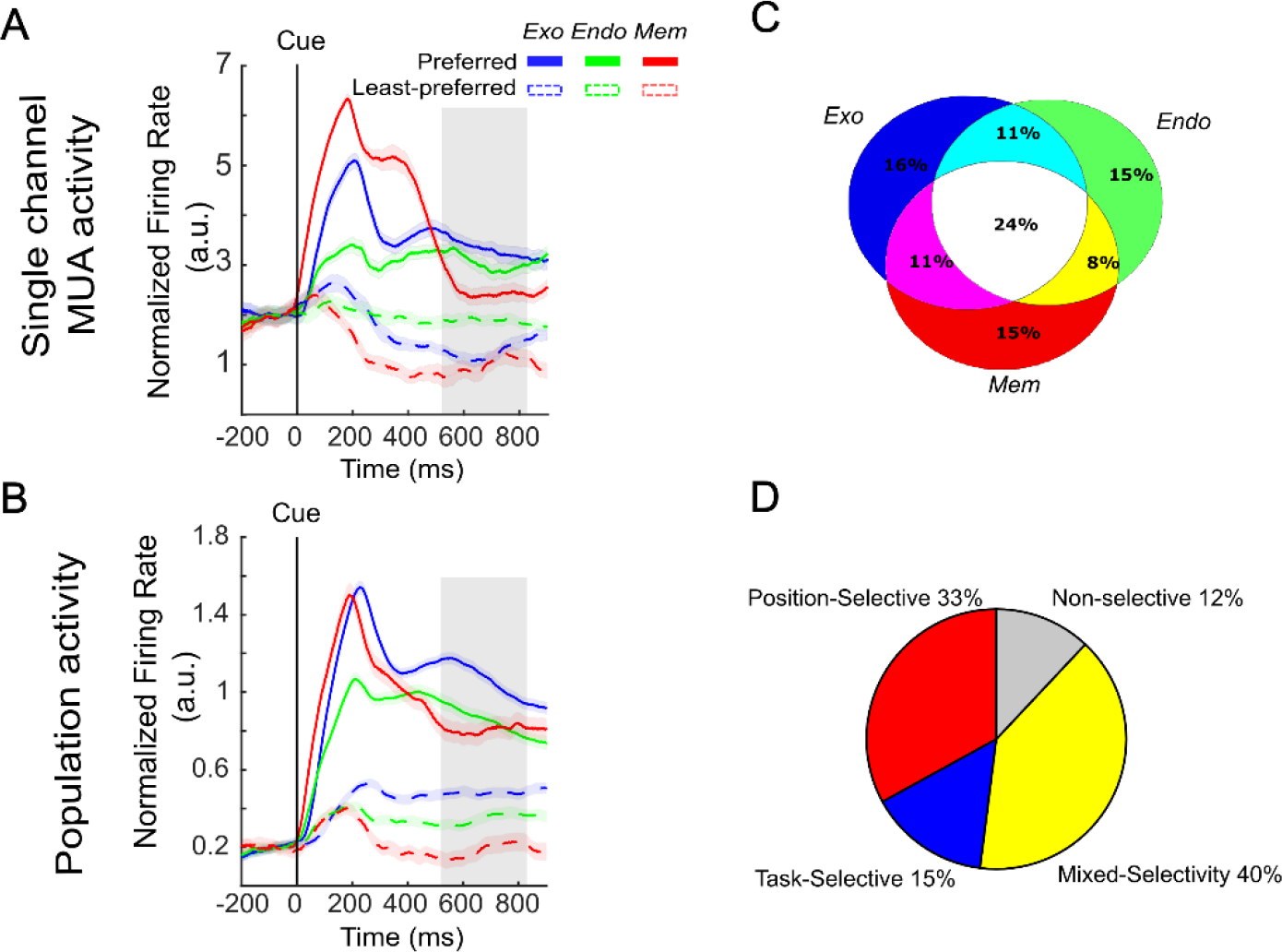
Mixed selectivity corresponds to the ability of a neuronal population to encode simultaneously independent sources of information: (A) Firing rate of the same MUA in the three tasks aligned on the cue (200 ms before the cue onset to 1000 ms after the cue onset) in exogenous task (blue), endogenous task (green) and memory guided saccade task (red) for preferred (full line) and non-preferred (dash line) position. (B) Mean firing rate of the same neuronal population in the three tasks aligned on cue presentation (200 ms before the cue onset to 1000 ms after the cue onset) in the exogenous task (blue), the endogenous task (green) and the memory guided saccade task (red) for preferred (full line) and non-preferred (dash line) position. (C) Venn diagram representing the proportion of cells showing a spatial tuning in each of the three tasks. (D) Pie chart corresponding to the proportion of neurons tuned to only position (red), task (blue) and both (mixed-selectivity; yellow). Non-selective neurons are plotted in grey.

Specifically, we observed that some cells were recruited during the performance of the three tasks during the delay period (Figure 2A). We measured the proportion of cells that expressed a significant spatial selectivity during the delay period of the three tasks. Results of this measure are represented in Figure 2C as a Venn diagram. Interestingly, we found that 24% of cells were tuned to the three different tasks, suggesting that there is a neural population encoding multiple cognitive processes (spatial attention & spatial working memory). In addition to the spatial selectivity in each of the three tasks, we observed that the activity of the neuron represented in the Figure 2A also varied as a function of the task. That is to say, we found neurons that showed mixed selectivity for both task and spatial information, as the one represented in this Figure. Therefore, we classified all recorded cells regarding either their selectivity toward position, task, or toward simultaneous position and task selectivity. Based on the statistical significance of the firing rate modulation for these two parameters (spatial selectivity and task selectivity), we found that 15% of cells were uniquely selective for the task, 33% were uniquely selective for position, and 40% showed a mixed selectivity for both parameters (Figure 2D). These results indicate that FEF encodes spatial information and the underlying cognitive processes recruited by each task through mixed-selectivity neurons.

### Task-related information accounts for FEF neuronal variability before cue presentation

In the previous section, we described that the averaged firing rate regime in response to the cue and during the delay epoch differed as a function of the type of task that monkeys were performing. We hypothesize that such different firing rate regimes as a function of the type of task come as a result of a context-dependent functional reorganization of the neural population. To test this hypothesis, we used dimensionality reduction to identify the low-dimensional representation of the neural recordings. Dimensionality reduction consists of projecting the high-dimensional neuronal recordings onto a low-dimensional space minimizing the information loss. To this end, we applied principal component analysis (PCA) to our data pooled along task category in a pre-cue time interval. At this time of the trial, task-related events soliciting the processing of spatial information have not been presented yet. Because trials from any given task are organized in blocks, at this time in the trial, the only prior information available to the monkey is task-related information. The projection of this pre-cue neuronal population data onto the principal components of the PCA showed mixing of task-information across the different components, as task-related neural variability was projected onto multiple PCA axes (Supplementary Figure 2). In the following we will use the term task-related information to refer to the information available in the neuronal population about task identity, i.e. about which task the monkey is engaged in.

Other methods of dimensionality reduction allow to label the explained variability by projecting specific sources of variance held in data onto each principal component. One of these methods is the demixed principal component analysis (dPCA)(Kobak et al., 2016). The objective of this method is to capture the majority of the variance in data in only few latent variables or components (as PCA does) but demixing the dependencies of the population activity on specific parameters by not constraining the components to be orthogonal. This method thus allows to directly interpret the impact of a related feature (e.g. task, position) on the neural population encoding. We applied dPCA on FEF MUA neuronal population activity in order to extract components related to task-related variability unmixed from other type of variability. To do this, we pooled data across task category (*Exo, Endo* and *Mem*). This analysis was firstly performed during the pre-cue period (200 ms before cue onset to cue onset) and will be later performed during the cue to target interval (500 ms to 0 ms before target onset) to unmix task-from sensory-specific variance.

The dPCA indicated that 90 % of the variance of the recorded brain activity prior to cue presentation was associated with task (Figure 3A). The overall variance explained by dPCA is similar to the variance explained by PCA (Figure 3B) indicating that dPCA reduces dimensionality of the data with a similar performance as PCA, thus minimizing the neuronal information loss. The three first dPCs (i.e. the two first task-related components (#1 and #2), and the first task-independent component (#3)) explained 80% of the variance, indicating that with only few components we recovered the majority of data variance. Therefore, population activity was accurately represented by the dPCA components. The angle between the two first components related to the task, did not significantly deviate from orthogonality (88°). This was confirmed by a second analysis performed on individual sessions (mean = 82°, std = 4°). This thus indicated that these two first components represented independent task-related information.

**Figure 3.**
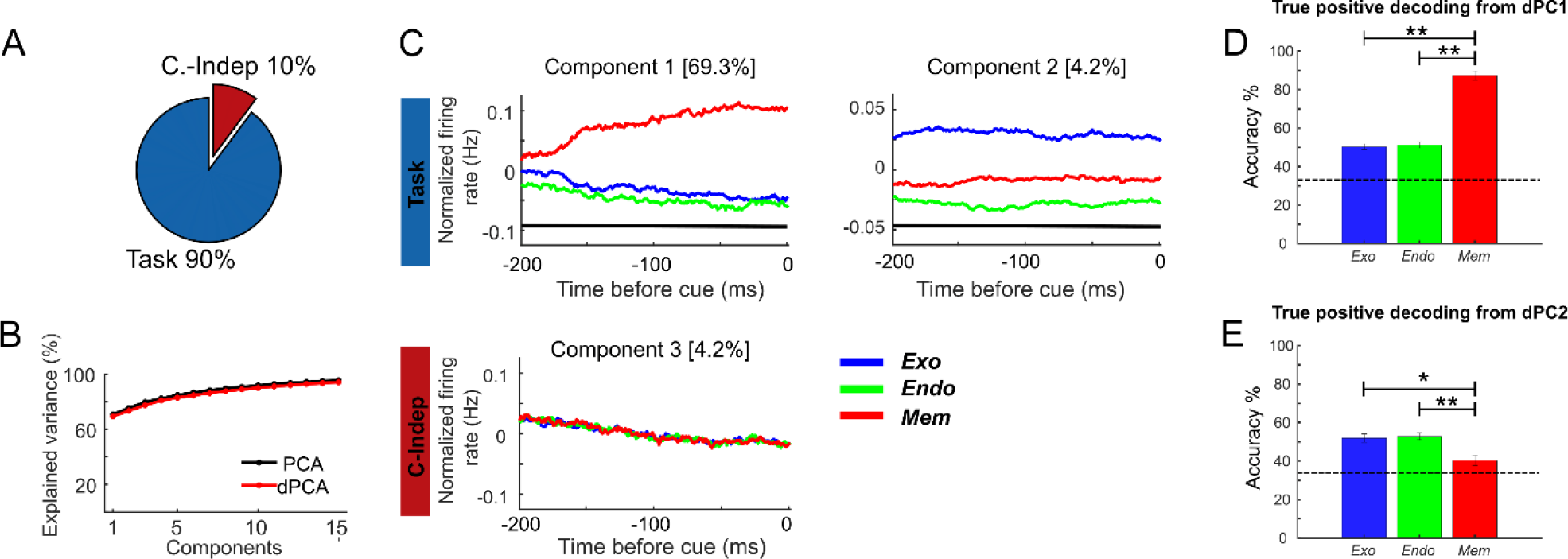
Demixed PCA unmixes task and independent sources related to variance. (A) Pie chart shows how the total signal variance is split among parameters: task (blue) and condition independent (red). (B) Cumulative variance explained by PCA (black) and dPCA (red). Demixed principal component show similar explained variance as PCA. (C) Demixed principal components. In each plot, normalized MUA firing rate averaged per task (in arbitrary units, exogenous (blue); endogenous (green) and memory-guided saccade (red) in the interval from -200 ms to 0 ms locked to the cue onset are projected onto the task-related dPCA components. Thick black lines show time intervals during which the task-information can be reliably extracted from single-trial activity (assessed against 95% C.I.). (D) True positive decoding rate for PC1 task-related. (E) True positive decoding rate for PC2 task-related. Dashed lines in (C) and (D) represent the 95% C.I.

Firing rates projected onto the first dPCA component associated to the task show a dissociation between the two attentional tasks and the working memory task (Figure 3C). To note, such difference cannot be interpreted as related to the response movement effector (hand response or saccade), since data was analysed during pre-cue activity (prior to motor preparation). The projection of the firing rates onto the second dPCA associated to the task show a dissociation between the exogenous (blue) and endogenous (green)s tasks suggesting a dissociation as a function of the attentional task-demands specifically within the attentional tasks. Supplementary Figure 3 shows, per each session, the projection of the dataset onto each of these components per task (trial-based). The third dPCA is associated with task-independent variance.

In order to assess whether the tuning of each of these two demixed components was statistically significant, we used these components as linear decoders to measure their ability to encode task-related information. To this end, we used cross-validation to measure time-dependent classification accuracy and shuffling procedure was used to assess the significance of the accuracy measure (see Methods for more details about decoding). Confusion matrix of each classification was extracted to analyse the true positive rates (TPR) of the classifier for each task. Figure 3D shows that for the first dPC, the TPR in classifying the memory task was 80% (iqr 7.3), while the TPR for both attentional tasks were significantly lower and not significantly different one from the other (2-way ANOVA: p = 1.0013e-15, *Exo vs Mem*: post-hoc Bonferroni test: p = 7.98e-15; *Endo vs Mem*: post-hoc Bonferroni test: p = 1.4e-14; *Exo vs Endo*: post-hoc Bonferroni p>0.05) suggesting a dissociation between attentional and working memory processes. In contrast, TPR in predicting the memory task using the dPC was lower than predicting both exogenous and endogenous attention (2-way ANOVA: p = 0.0039, E*xo vs Mem*: post-hoc Bonferroni: p = 0.0137; *endo vs mem*: post-hoc Bonferroni: p = 0.0076; *Exo vs Endo*: post-hoc Bonferroni: p>0.05) suggesting a dissociation in the way the attentional processes (exogenous vs endogenous) were implemented.

### Task-related and spatial information segregate in orthogonal demixed components

Mixed selectivity corresponds to the ability of neurons to simultaneously encode different sources of information (Fusi et al., 2016; Rigotti et al., 2013). The aim of the present analysis was to first investigate whether these different sources of information could be assigned to separate principal components and then to quantify the level of their interaction. Here, we were interested in the task and target spatial information, thus, we will focused on the delay period between the cue and the target in order to include spatial information, as opposed to the pre-cue interval that had been selected in our previous analysis in which there was no spatial information available yet. We applied dPCA on neuronal population activity during the delay epoch, respectively organized by position independently of task, and by task independent on position, in the time interval corresponding to cue to target interval (500 ms to 0 ms before target onset) (Figure 4). Demixed PCA split variance in position-related variance (52% of the explained variance), task-related variance (26% of the explained variance); and interaction between task information and spatial information (20 % of the explained variance) (Figure 4A). The rest of the variance (2%) related to independent sources of information. The overall variance explained by dPCA was similar to that explained by PCA, both accounting for 70% of the variance by the five first components. As for the pre-cue period (see previous section), we found that the first task-related dPC did not significantly deviate from orthogonality relative to the second task-related dPC during the cue-to-target interval (87.9°), and both of these did not significantly deviate from orthogonality relative to the first position-related dPC (81.6°; respectively, 89.8°). We reproduced these observations when performing this analysis on the independent sessions (dPC1-task vs dPC2-task: mean = 83°, std = 5°; dPC1-pos vs dPC1-task: mean = 74°, std = 8°; dPC1-pos vs dPC2-task: mean = 83°, std = 7°). Therefore, population activity was accurately represented by different orthogonal dPCA components (Figure 4B). Similarly to what we found in the previous section, the two first components associated to the task showed respectively a dissociation between attentional task-related processes and working memory task-related process (component 3), and a trend of separation between the two attentional tasks (component 4). In addition, the first component associated to position exhibited a dissociation of each position corresponding to the four expected target locations, whereas the projection onto the second spatial-related component showed two different states possibly corresponding to the segregation of information across hemifields and recording probes (i.e., probes respectively located into right or left FEF) (supplementary Figure 4). Last, the interaction component (Figure 4C low panel) describes the non-linear mixed selectivity between information related to task and position. We used this interaction component to decode trials based on the cued position and task. Figure 4D shows the average confusion matrix of this decoder across all sessions. We found that true positive rates for each of the twelve classes were significantly above the 95% C.I. (Supplementary Figure 4C). Overall, this indicates that part of the spatial information represented in the FEF neuronal population is independent (orthogonal) to the task-related information, while another part of this spatial information interacts with the task-related information.

**Figure 4.**
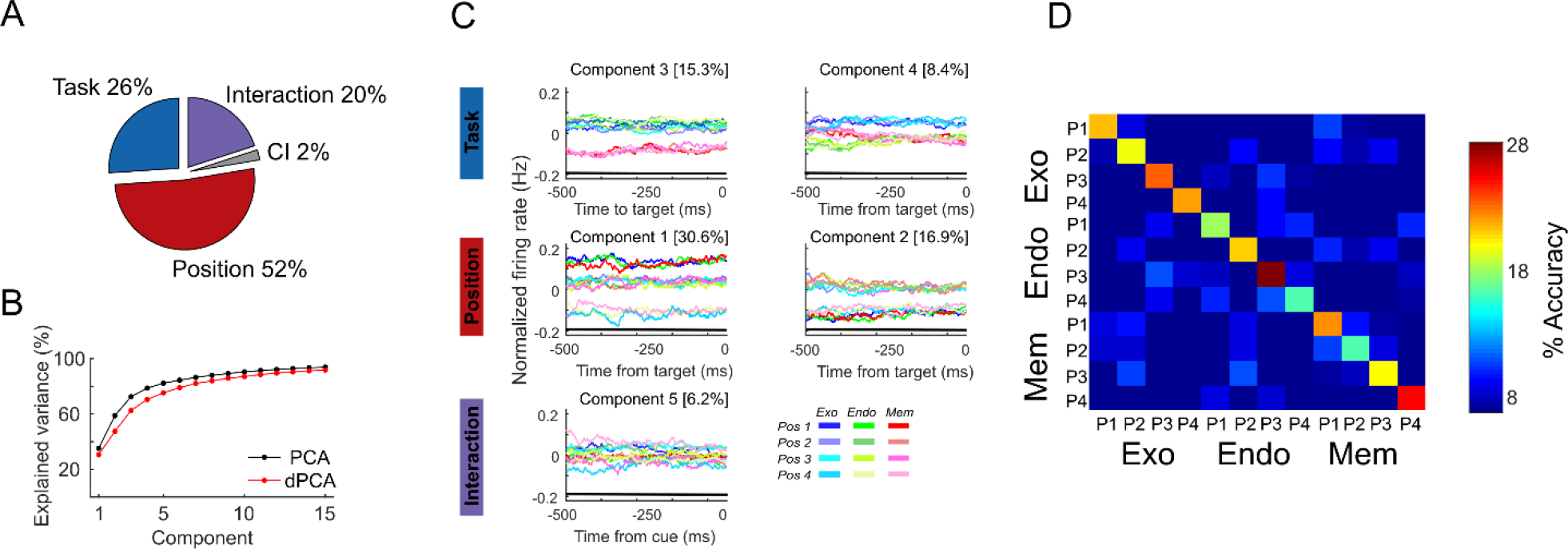
Demixed PCA unmixes task and spatial sources related to variance. (A) Pie chart shows how the total signal variance is split among parameters: Task (blue), Position (red), Interaction (purple) and Condition independent (grey). (B) Cumulative variance explained by PCA (black) and dPCA (red). Demixed principal components show similar explain variance as PCA. (C) Demixed principal components. In each plot, MUA firing rate averaged per tasks (exogenous (blue shades); endogenous (green shades) and memory guided saccade (red shades)) and cued position in each task, in the interval from -500 ms to 0 ms locked to target onset are projected onto the task-related, position-related and interaction dPCA components. Thick black lines show time intervals during which the parameter information can be reliably extracted from single-trial activity (assessed against 95% C.I.). (D) Confusion matrix of decoding task and position using the demixed interaction components, averaged across all sessions. All elements along the diagonal are significantly above the 95% C.I.

## Discussion

In this manuscript, we addressed the following two questions. First, we asked whether information about the ongoing task identity could be extracted from the FEF. Second, we asked whether the encoding of the spatial information in the FEF necessary for the performance of a task depended on task-identity, that is to say, depended on the cognitive processes underlying the task requirements. To address this question, we analysed MUA recorded intracranially from the FEF, bilaterally, while two monkeys performed three cognitive tasks in the same recording session. These three tasks presented a very similar structure (the onset of a cue was followed by a target stimuli). However, they recruited different underlying cognitive mechanisms necessary for their performance: endogenous spatial attention, exogenous spatial attention, or spatial working memory. We found that, at the neural level, the dynamics of the activity of the FEF cells when their preferred position was cued was different across tasks, suggesting a different encoding of the sensory information as a function of the task being performed. Additionally, we found a remarkable population of mixed-selectivity cells that showed a simultaneous encoding of position and task-identity. In order to understand how this information was organized at the population level, we applied a supervised dimensional reduction method (dPCA, (Kobak et al., 2016; Machens, 2010)) which unmixed the neuronal variability attributed to different sources of information onto different components. We found that the FEF encodes task identity in two sources of task-related variability (independently from the processing of sensory information) that segregate in orthogonal components that conforms a two-dimensional space, with one component dissociating spatial working memory vs spatial attention processes, and the other component dissociating specifically both spatial attentional processes (endogenous vs exogenous). Additionally, we observed that the neuronal population encodes task-identity and trial-relevant spatial information in a unique interaction component that can be decoded using a single linear readout. This is a hallmark of a high dimensional representation of sensory information. All in all, we show that the FEF have a flexible functional architecture through which it encodes differently the same spatial information as a function of the task demands and underlying cognitive processes.

Prior evidence showed that FEF encodes spatial information during the performance of attentional tasks (Amengual et al., 2022; Armstrong et al., 2009; Bruce and Goldberg, 1985; Di Bello et al., 2021; Goldberg and Segraves, 1987; Paneri and Gregoriou, 2017) and working memory tasks (Baddeley, 2012; Bahmani et al., 2019; Funahashi and Kubota, 1994; Noudoost et al., 2021, 2021; Sapountzis et al., 2022). Interestingly, Bahmani and co-workers (Bahmani et al., 2019) described a functional overlap between these two cognitive processes in the FEF, and how the coincident impairment of working memory and attention might be associated with disruptions of dopamine signals in this area. Taken together, these previous findings suggest that the FEF is able to encode simultaneously different sources of information, related to sensory or cognitive processes associated with specific tasks. In the present study, we confirm that the FEF neuronal population is able to treat the same sensory information differently as function of the context set by task identity. This is achieved thanks to a high dimensional representation of the neuronal population which endows the FEF with a high degree of flexibility. The neural mechanisms underlying this coding strategy are discussed next.

We propose that such flexible encoding ability of the FEF results from the activity of mixed-selectivity cells that are present in the recorded neuronal population, in agreement with previous studies (Amengual et al., 2022; Fusi et al., 2016; Mendoza-Halliday and Martinez-Trujillo, 2017; Rigotti et al., 2013). In our data, we found that 40% of the recorded neurons show simultaneous tuning for position and task-identity. In addition, more than half of the recorded neurons are involved in encoding spatial information in at least two of the tasks performed, and 24% of the recorded cells exhibit spatial tuning in all of the three tasks. Such a rich functional configuration of this area leads to its ability to encode sensory information differently as function of the context (aka, task-identity). As a result, we applied a dimensionality reduction method (demixed PCA, dPCA, (Kobak et al., 2016; Machens, 2010)) on the recorded neuronal population in order to split the variance in the data in specific components, which variance is associated to either the sensory information, task-identity and their interaction. To note, this method is different to the PCA, which decomposes the variance in different components without considering its functional source.

Applying the dPCA during a time-interval prior to cue onset (i.e. corresponding to baseline activity prior to the presentation of the specific ongoing trial spatial requirements, thus reflecting task-related neuronal variability independently from sensory processes, as trials were presented in blocks), we found a two-dimensional subspace of orthogonal components the activity of which was linked to task-identity. One of these components dissociated the two main cognitive processes associated with these three tasks (spatial attention variants and spatial working memory), while the second component specifically dissociated the two types of attention in the exogenous task (in which the cue was presented peripherally), and attention in the endogenous task (in which the cue was presented centrally). These results are in line with prior studies (Bahmani et al., 2019; Panichello and Buschman, 2021), and showed the ability of this area to modulate its patterns of neuronal responses as a function of underlying cognitive process.

A similar pattern of activity was observed after the presentation of the cue, prior to target presentation. That is to say, when dPCA was applied at the interval of time corresponding to the delay period (therefore, when the position information is maintained), we observed the same two-dimensional orthogonal components relative to task identity. This suggests strongly a stable state of the underlying cognitive information encoded by the neuronal population in time, similarly to what has been described in other PFC recordings (Amengual et al., 2022; Cowley et al., 2020; Oby et al., 2019). Additionally, a two-dimensional space composed of orthogonal components related to the sensory information was observed specifically during the delay period. This suggests that FEF simultaneously encodes sensory information and task identity by using different components. Importantly, we found another component related to the interaction between task-identity and the cued position. When we trained a decoder to simultaneously decode task-identity and position information using the firing rates projected onto this interaction component, we found that the accuracy of this classifier was above chance level, indicating that all possible information instances (four cued positions in each task) were decoded. This represents a marker of a high dimensional representation of information in the neuronal population, and it suggests that the FEF specifically encodes the sensory information as a function of the ongoing task, hence as a function of the cognitive process required by the specific task. These results extended those reported by Astrand and colleagues (Astrand et al., 2015), where they showed a different population dynamics and code stability as a function of the source of the information and task. Specifically, they showed, during attentional task dissociating attention orientation from the colour and the position of the instruction cues, a stable time-resolved accuracy of decoding spatial attention in attentional tasks, in contrast to the dynamic encoding of colour and stimulus position. Other studies from the same team showed a different encoding dynamics of sensory information as a function of whether attention was cued endogenously (centrally) or exogenously (peripherally) (Astrand et al., 2016). All in all, our results reinforce previous studies suggesting that the dynamics of the encoding of sensory information might be different based on the underlying cognitive process recruited by the task.

Whether such task-dependent sensory encoding that we have observed in the FEF is inherent of its own functional architecture or a consequence of the interaction between its activity and the activity of other brain regions at a network level is something that, unfortunately, we cannot tackle in this study, although is worthy to discuss. Particularly, we have focused on intracranial recordings on the FEF, a key region in the PFC involved in top-down attentional process (Corbetta and Shulman, 2002). However, it is well known that the FEF, given its rich anatomical connectivity, is a functional hub showing efferent and afferent connections to other brain areas such as the dorsolateral prefrontal cortex, the cingulate cortex, the parietal cortex and the superior colliculus, amongst others (Vernet et al., 2014 for review). In the attention domain, it is known that there is a dual information flow between the FEF and the lateral intra-parietal area (LIP) during the attention-task performance (Ibos et al., 2013). Other studies have found an active functional role of the LIP during visual working memory tasks (Johnston et al., 2010). Therefore, one possibility would be that the functional network linking the FEF and the LIP might have a differential information flow regarding the sensory information, which would impact on the way the FEF encodes top-down sensory signals. Other studies abovementioned have described the functional role of the prefrontal dopaminergic cells in both attention and working memory (Bahmani et al., 2019). Indeed, dopamine has been shown as a common modulator of attention and working memory, and its imbalance in patients affected by Parkinson’s disease have been accompanied by a clear impairment in both cognitive processes. Therefore, another possibility could be that task-identity activity described in the demixed components might be associated with a differential modulation of the activity of these prefrontal dopaminergic cells driven by the task. Another possibility yet would be that these effects of the modulation of the dopaminergic system in the FEF might also impact on the ability to decode the sensory information, reflecting a different encoding strategy of this information based on task-identity. In order to address these hypotheses, further pharmacologic studies based on the dopaminergic prefrontal cortex regulation should be performed.

In conclusion, we show the potential capacities of the FEF in encoding sensory information, and how this information encoding is performed differently as a function of the cognitive demands of the underlying task. We believe that these results provide novel insights on how the FEF organizes, from a computational perspective, attention and working memory information and recruits them in the context of specific tasks, and how potential pharmacological interventions can be targeted to improve these cognitive processes in patients affected by neurological diseases. Moreover, this knowledge leads to an improvement in the design of cognitive brain computer interface field (cBCI), opening the venue to the manipulation of the activity not only related to sensory or motor processes, but also related with a specific cognitive modality (Astrand et al., 2014; Loriette et al., 2022, 2021).

## Author contributions

Conceptualization: C.W., J.A. and S.B.H.; Data Curation: E.A., A.M.; Formal Analysis: A.M., E.A., J.A.; Funding Acquisition: S.B.H.; Investigation: E.A., C.W., S.B.H.; Methodology: J.A., S.B.H.; Resources: A.M., E.A., C.G., J.A., S.B.H.; Supervision: J.A., S.B.H.; Validation: J.A., S.B.H.; Visualisation: A.M., J.A.; Writing-Original draft: A.M., J.A. and S.B.H.; Writing – Review and Editing, A.M., E.A., C.W., C.G., J.A. and S.B.H.

## Acknowledgements

We thank engineer Serge Pinède for technical support, Jean-Luc Charieau and Fabrice Hérant for animal care, and Johan Pacquit, Sylvain Maurin and Thomas Perret for informatics assistance. All procedures were approved by the local animal care committee (C2EA42-13-02-0401-01) in compliance with the European Community Council, Directive 2012/63/UE on Animal Care. J.A. is now affiliated to *Instituto de Investigación Sanitaria La Fe, Plataforma de Big Data, IA y Bioestadística, 46026, Valencia, Spain*; C.G. is now affiliated to *Laboratory in Sensory Physiology, Otto Von Guericke University, 39120, Magdeburg, Germany;* E.A. is now affiliated to *School of Innovation, Design, and Engineering, Mälardalen University, IDT, 721 23 Västerås, Sweden;* C.W. is now affiliated to *UMR 1253, iBrain, Université de Tours, Inserm, Tours, France*.

## Data and code availability

All data reported in this paper will be shared by the lead contact upon request. All original codes will be made available at time of publication through a public repository.

## Supplementary Figures

**Supplementary Figure 1.**
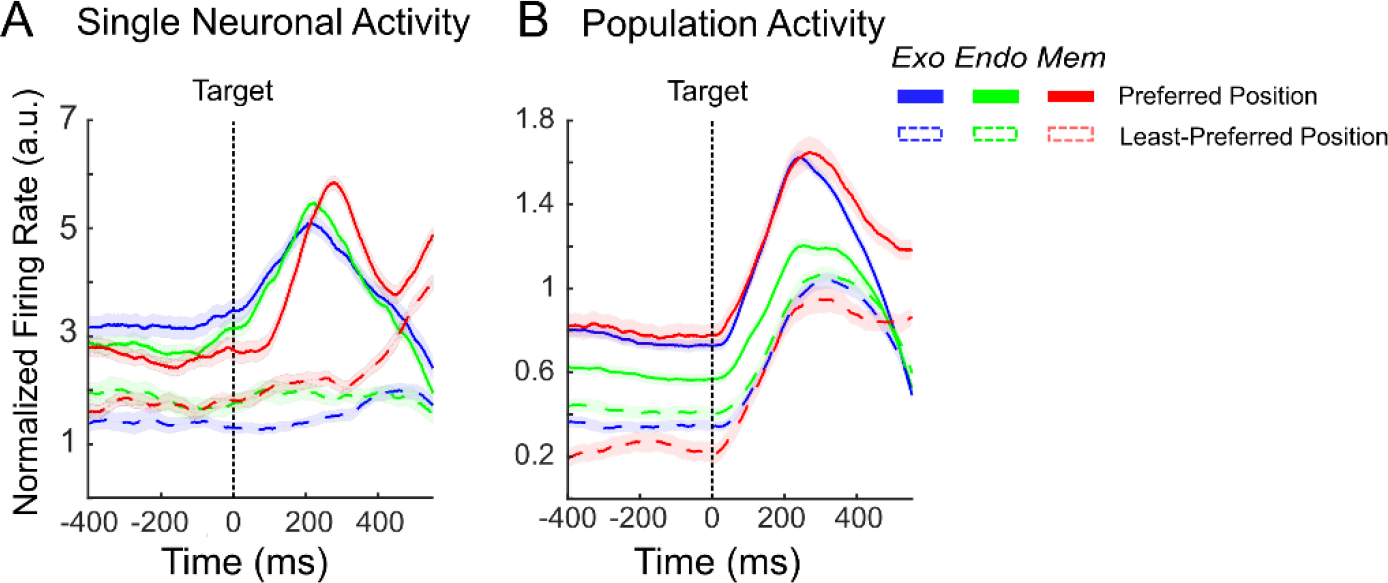
(A) Firing rate of a neuron (same as in Figure 1) aligned to target onset (400 ms before target onset to 600 ms after target onset) averaged across trials of the exogenous attention task (blue), the endogenous attention task (green) and the memory guided saccade task (red) for preferred (full line) and least-preferred (dash line) positions. (B) Mean firing rate across the neuronal population (MUA) aligned on the target (-400 ms before target onset to 600 ms after target onset) as a function of tasks and positions (same as in A).

**Supplementary Figure 2.**
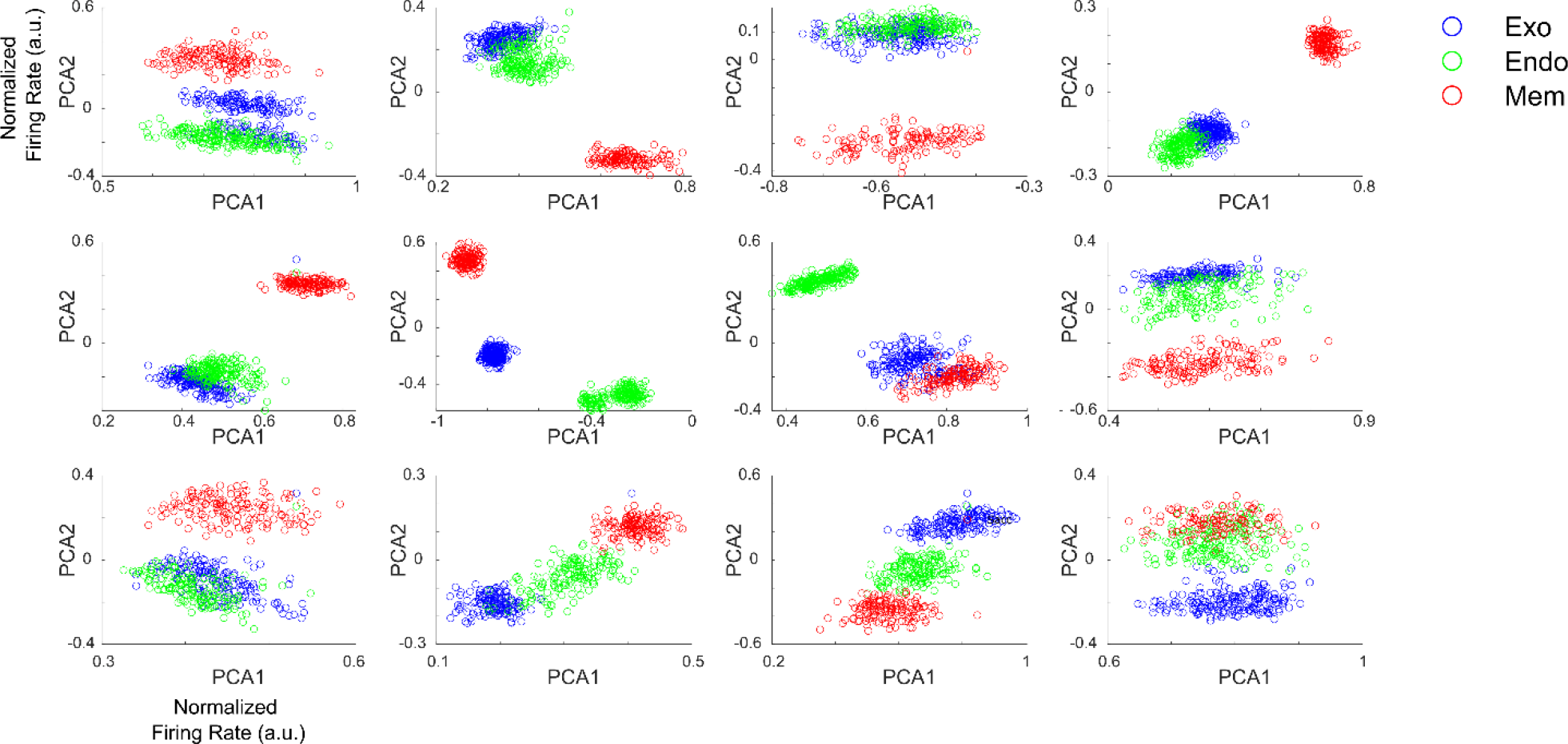
Projection of MUA onto the first and second principal components extracted with a PCA analysis. Each point corresponds to the projection of the activity in each trial (averaged in the time interval -300 to - 100ms pre Cue, each circle thus corresponds to one trial) in each task (Exogenous task, blue; Endogenous task; green; Memory saccade task, red). Each plot corresponds to a different session. Plots 1 to 7 are from monkey 1 and plots 8 to 12 are from monkey 2.

**Supplementary Figure 3.**
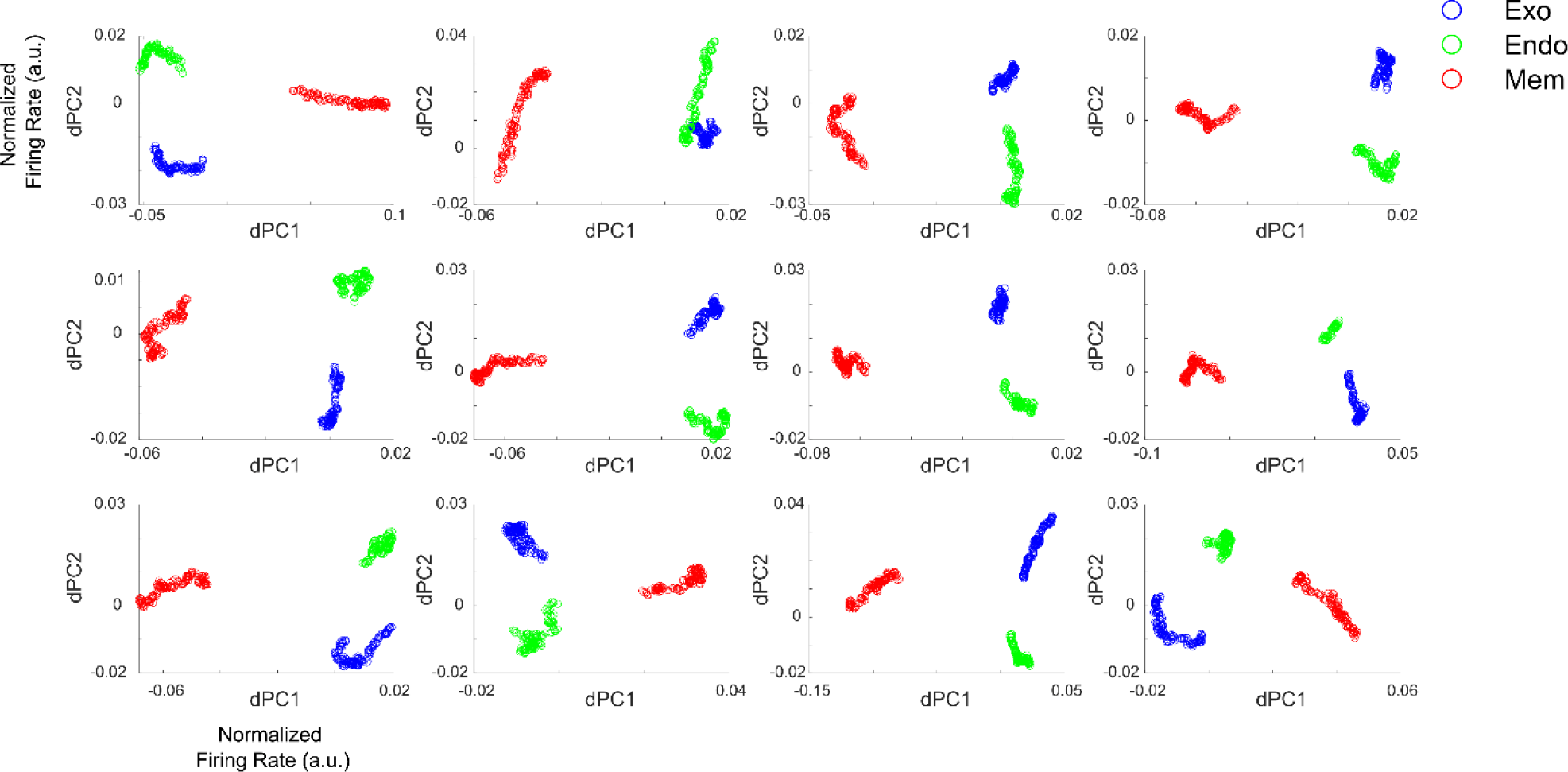
Projection of MUA onto the first and second demixed principal components extracted with a dPCA analysis. Each point corresponds to the projection of the activity in each trial (averaged in the time interval - 300 to -100ms pre-Cue, each circle thus corresponds to one trial) in each task (Exogenous task, blue; Endogenous task; green; Memory saccade task, red). Each plot corresponds to a different session. Plots 1 to 7 are from monkey 1 and plots 8 to 12 are from monkey 2.

**Supplementary Figure 4.**
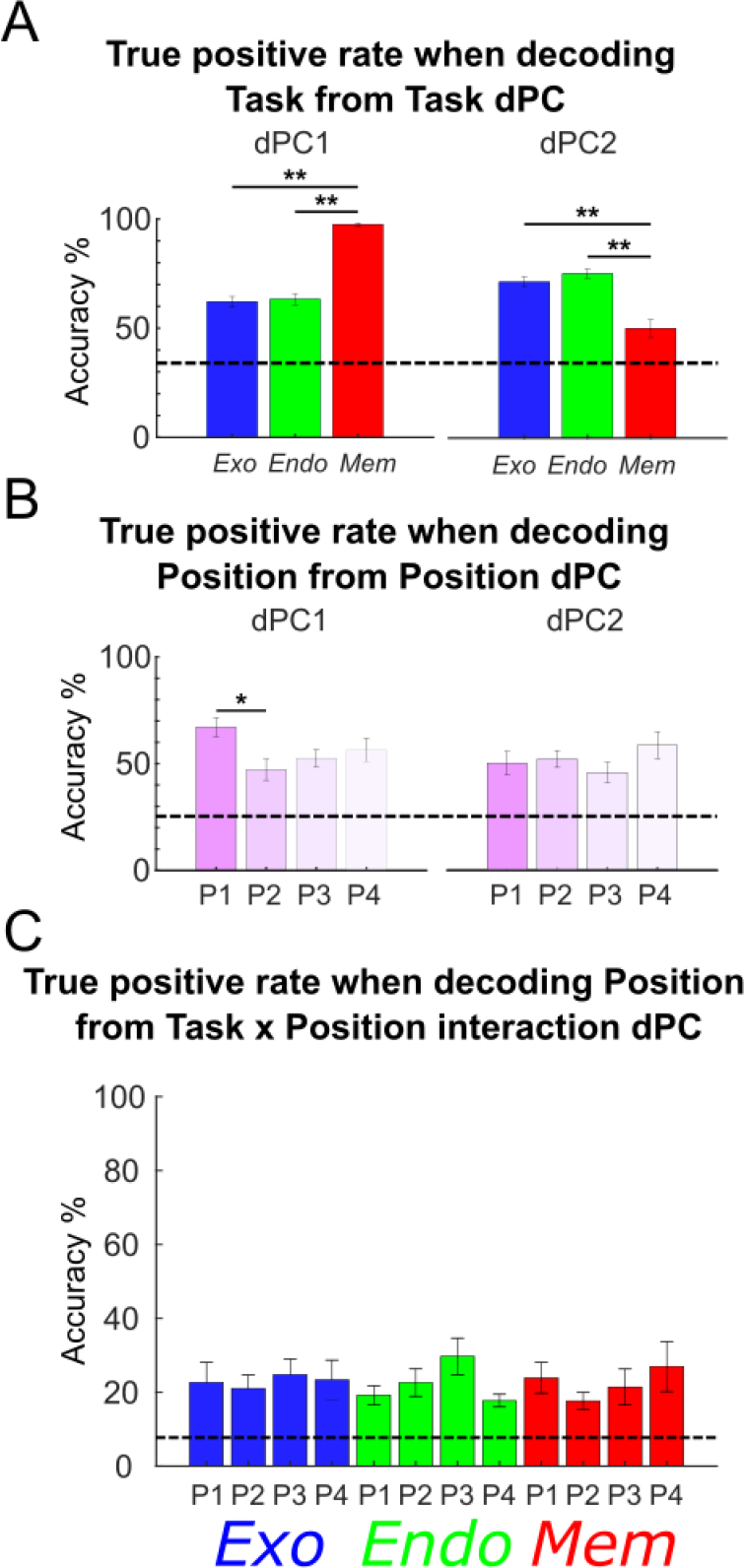
(A) Barplot corresponding to the true positive rates obtained using the first task-related demixed principal components as a linear classifier to classify trials by task (Endogenous attention, blue; Exogenous attention, green; Memory saccade, red) (2-way ANOVA test dPC1: p = 4.5e-15: exo vs endo: post-hoc Bonferroni: p = >0.05; exo vs mem: post-hoc Bonferroni: p = 3.3e-14; endo vs mem: post-hoc Bonferroni: p = 6..6e-14; 2-way ANOVA test dPC2: p = 7.23e-9, exo vs endo: post-hoc Bonferroni: p>0.05; exo vs mem: post-hoc Bonferroni: p = 2.51e-7; endo vs mem: post-hoc Bonferroni: p = 1.54e-8). (B) Barplot corresponding to the true positive rates obtained using the first position-related demixed principal components as a linear classifier to classify trials by position 2-way ANOVA test dPC1: p = 0.0277; pos1 vs pos2: post-hoc Bonferroni: p = 0.0235; other positions comparisons showed no significant difference, p>0.05); 2-way ANOVA test dPC2: p = 0.4198. (C) Barplot corresponding to the true positive rates obtained using the first task x position interaction demixed principal component as a linear classifier to classify trials by position in each task (Friedman test: p = 0.8539, Interaction task x position showed no significant difference, p>0.05).

